# A binding site for the antibiotic GE81112 in the ribosomal mRNA channel

**DOI:** 10.1101/2024.09.26.614503

**Authors:** Andreas Schedlbauer, Xu Han, Wouter van Bakel, Tatsuya Kaminishi, Borja Ochoa-Lizarralde, Idoia Iturrioz, Retina Çapuni, Ransford Parry, Ronny Zegarra, David Gil-Carton, Jorge P. López-Alonso, Kristina Barragan Sanz, Letizia Brandi, Claudio O. Gualerzi, Paola Fucini, Sean R. Connell

## Abstract

The initiation phase is the rate-limiting step of protein synthesis (translation) and is finely regulated, making it an important drug target. In bacteria, initiation is guided by three initiation factors and involves positioning the start site on the messenger RNA within the P-site on the small ribosomal subunit (30S), where it is decoded by the initiator tRNA. This process can be efficiently inhibited by GE81112, a natural hydrophilic, noncyclic, nonribosomal tetrapeptide. It is found in nature in three structural variants (A, B and B1 with molecular masses of 643-658 Da). Previous biochemical and structural characterisation of GE81112 indicates that the primary mechanism of action of this antibiotic is to (1) prevent the initiator tRNA from binding correctly to the P-site and (2) block conformational rearrangements in initiation factor IF3, resulting in an *unlocked* 30S pre*IC* state. In this study, using cryoEM, we have determined the binding site of GE81112 in initiation complexes (3.2-3.7Å) and on empty ribosomes (2.09 Å). This binding site is within the mRNA channel (E-site) but remote from the binding site of the initiation factors and initiator tRNA. This suggests that it acts allosterically to prevent the initiator tRNA from being locked into place. The binding mode is consistent with previous biochemical studies and recent work identifying the key pharmacophores of GE81112.

## INTRODUCTION

In all organisms, the initiation phase is the rate-limiting step of protein synthesis (translation) and is subject to fine-tuning via posttranscriptional regulatory mechanisms (Duval et al., 2015; Gualerzi and Pon, 2015). As such, the initiation step in protein synthesis is an important drug target (Brandi et al., 2007). The initiation phase is a multistep, dynamic process that begins with the formation of a 30S pre-initiation complex (pre*IC*). In the pre*IC,* the mRNA, tRNA, and three protein factors, termed initiation factors (IFs: IF1, IF2, and IF3), assemble on the small ribosomal subunit (30S subunit) to recognise the mRNA start codon (Gualerzi and Pon, 2015). In this state, the tRNA is liably bound (Milón et al., 2012), and therefore it is referred to as an unlocked pre*IC*. This unlocked complex undergoes a conformational change where the carboxy-terminal domain (CTD) of IF3, positioned at the top of h44 in the pre*IC*, moves to a second position on h44 (Julián et al., 2011; Hussain et al., 2016; López-Alonso et al., 2017). This movement allows the tRNA to be fully accommodated in the P-site, locking the fMet-tRNA in place and forming a 30S initiation complex (30*IC*). The resulting 30*IC* joins the large ribosomal subunit (LSU, the 50S), releasing IF1 and IF3 to form the 70S ribosomal initiation complex (70S*IC*).

GE81112 (GE8) is a specific inhibitor of this initiation process and blocks the locking step that marks the transition from the pre*IC* to the *IC* (Brandi et al., 2006a; Fabbretti et al., 2016). GE81112 is a peptide antibiotic formed by four non-proteinogenic L-amino acids, referred to as AA1-4. When purified from the producing *Streptomyces* species, GE81112 is a mixture of three congeners, termed GE81112A, GE81112B, and GE81112B1 (643-658 Da)(Brandi et al., 2006b). Owing to its novel chemical scaffold and potential as a lead molecule for antibiotic development, several enzymatic and chemical synthesis pathways have been established for GE81112 (Jürjens et al., 2018; Zwick et al., 2019, 2021; Fayad et al., 2023). This has allowed structure-activity relationship (SAR) studies to define GE81112 key pharmacophores (Jürjens et al., 2018; Zwick et al., 2021). For example, the work of Zwick *et al*. (Zwick et al., 2021) highlights the importance of AA1 and AA4 for antibacterial activity. The Bauer group has also performed extensive pharmacokinetic profiling on GE81112A, finding several limitations but observing that GE81112 “could represent a promising starting point for chemical modifications and optimisation programs” (Schuler et al., 2023). Such programs would greatly benefit from improved structural data (Fabbretti et al., 2016; López-Alonso et al., 2017) describing the interaction of GE81112 with its ribosomal target. One limitation of GE81112 is its dependence on the peptidyl transporter oligopeptide permease (OPP) for entry into the cell (Maio et al., 2016). Because of this, GE81112 shows limited *in vitro* activity in rich media as it is outcompeted by peptides in the media for transport into the cell (Maio et al., 2016; Schuler et al., 2023). Moreover, as the OPP system is not essential for bacterial growth in laboratory conditions, mutations that inactivate OPP result in resistance to GE81112 (Maio et al., 2016). Modifying the GE81112 scaffold to overcome this dependence could enhance GE81112’s potential.

## RESULTS

### Structure Overview

To understand the interaction of GE81112 with the *E. coli* ribosome, *in vitro* reactions mimicking the initiation process (*E. coli* 30S subunits, fMet-tRNA, mRNA, IF1, IF2, IF3) were set up in the presence of GE81112 (see Material and Methods). The reaction conditions were established in previous FRET and cryoEM experiments, which showed that GE81112 traps the 30S in a preinitiation state (pre*IC*) (Fabbretti et al., 2016; López-Alonso et al., 2017). Under these conditions, two independent samples were prepared and used in single-particle cryoEM experiments (**Supplemental Figures S1 and S2**). The dataset obtained from the first sample (**Figure 1**, DATASET 1) showed density for all three initiation factors, albeit at a relatively low resolution (3.8 Å global resolution, complex 1: 30S+GE8+fMet-tRNA+IF1/IF2/IF3, **Supplemental Figures S1**). In comparison, the cryoEM maps from the second sample/dataset had a much weaker density for IF2 but showed higher resolution (**Supplemental Figure S2)**. plex 2: 30S+GE81112+fMet-tRNA+IF1/IF3 (3.3 Å consensus refinement; 3.2 Å Body; 3.2 Å Head), complex 3: 30S+GE81112+fMet-tRNA (3.3 Å consensus refinement; 3.3 Å Body; 3.7 Å Head), complex 4: 30S+GE81112 (3.1 Å consensus; 3.1 Å body; 3.1 Å head). In complex 1, where IF3 and fMet-tRNA are present, the C-terminal domain of IF3 (IF3-CTD) is seen bound to the top of h44 (pos1) and the fMet-tRNA is in a pre-accommodated position when compared to the initiation complexes of Hussain *et al*. (**Supplemental Figure S5**)(Julián et al., 2011; Hussain et al., 2016). This indicates the initiation reaction was stalled in an early initiation state, suggesting that GE is actively stalling the assembly process as expected (López-Alonso et al., 2017). In complexes 1-4, lower local resolution (**Supplemental Figure S1** and **S2)** was observed for the fMet-tRNA and initiation factors, indicating that in the pre*IC*, these elements are compositionally or conformationally dynamic. This compositional and conformational heterogeneity is consistent with the previous cryoEM studies on the 30SIC, which show multiple states and large-scale movements, particularly in the IF3-CTD (Hussain et al., 2016). For discussing the position of the IFs/tRNA and mRNA, we refer primarily to complex C as in dataset 1, the cryoEM map for the head region was interpretable without needing multibody refinements; Dataset 2 required the use of multibody refinements and the regions at the interface of the body, particularly the mRNA/tRNA, are difficult to interpret. Overall, the resolution of the cryoEM maps was sufficient to model the components of the 30S at a residue level. In contrast, the lower resolution (**Supplemental Figure S1** and **S2**) in the density of the initiation factors allowed for accurate modelling of the backbone at the secondary structure level.

**Figure 1:**
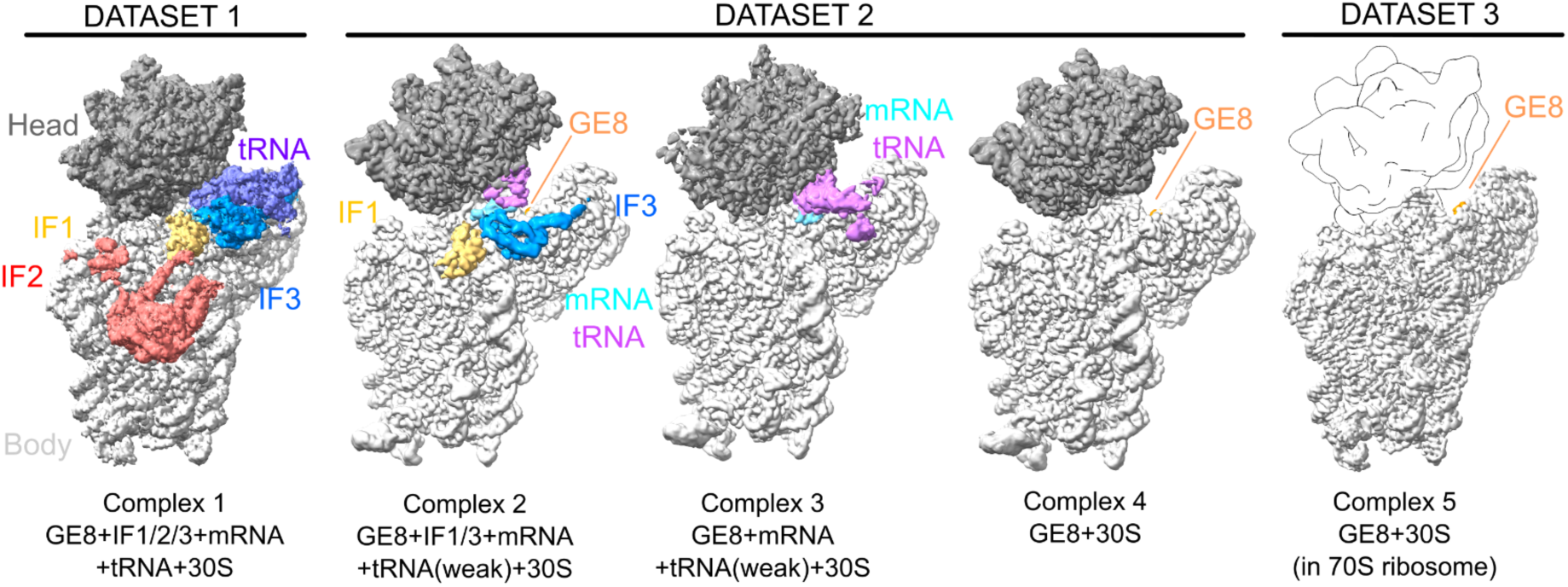
Cryo-EM Structure Overview. CryoEM maps segmented and colored to highlight density for the initiation factors, mRNA, fMet–tRNA, GE8, and 30S head/body domains. The cryoEM map in dataset 1 results from a refinement of the entire 30S subunit, dataset 2 from a multibody refinement where the head and body regions were refined separately, and dataset 3 from a refinement of the 30S body region after density corresponding to the 30S head and 50S subunit were subtracted. The maps shown are sharpened with deepEMhancer (Sanchez-Garcia et al., 2021) and were segmented using masks generated from the models. GE81112 is present in all maps but sometimes is obscured by density for other ligands. In dataset 3, a trace of the 30S head (based on a PDB model) is shown for clarity.

In addition to the initiation complexes, a third sample (**Figure 1**, DATASET 3) was prepared that included only GE81112 bound to a 70S ribosomal particle (**Supplemental Figure 3**) to determine a high-resolution model for the GE81112 binding pocket. Single particle cryoEM on this complex yielded a final cryoEM map for the 30S subunit body with an overall resolution of 2.09 Å (consensus 70S map: 1.9 Å; **Figure 1**). This high-quality map allowed the accurate placement of GE81112 in the map and an analysis of its interaction with the 30S subunit. Note this complex was prepared on a 70S ribosome to facilitate improved resolution; it is not meant to imply GE81112 targets 70S functions as, in fact, prior characterisation indicates it is an initiation targeting antibiotic (Brandi et al., 2006a).

### GE81112 binds to a pocket formed by h23, h24 and h45 in the mRNA channel

Previously, using X-ray crystallography and *Thermus thermophilus* 30S ribosome, we observed density consistent with GE81112 being bound close to the bent ASL of the P-site tRNA, represented in the crystal by helix 6 (h6) of a symmetry-related 30S subunit (Fabbretti et al., 2016). This binding site is distinct from the one we observe here, using cryoEM, on a bonafide *in vitro* assembled *E. coli* 30S initiation complexes and on *E. coli* 70S ribosomes. Here, GE81112 is seen bound in a pocket that overlaps with the E-site mRNA, formed by the grove of h24 (U788-G791, A794-C796), h23 (G693), h45 (U1506) and the CTD of r-protein S11 (**Figure 2A-C**). As seen in **Figure 2B**, the cryoEM density accommodates all atoms of GE81112, and in particular, the high-resolution (2.09 Å) complex 5 map, when sharpened, allows the orientation of the 3-hydroxy-L-pipecolic acid ring to be reliably determined; in complexes 1-4 this was ambiguous. Moreover, as mentioned, GE81112 purified from *Streptomyces spp*. (as used in our experiments) is a mixture of three congeners, termed GE81112A, GE81112B, and GE81112B1, with slight chemical differences in AA2 and AA3 where, as depicted in **Figure 2F**, either an oxygen or an NH group are present in AA2 at position ε, while an hydrogen or an NH2 group is present at position 2 in AA3. In this respect, although the complex 5 cryoEM map has a high local resolution in the GE81112 binding pocket (**Supplemental Figure 3**), it does not indicate the presence of a specific congener, considering we use a mixture of all three to prepare the complex. For example, the map suggests the presence of an NH2 group at position “R”, present only in congeners B and B1 (**Figure 2G**), but it does not exclude the presence of congener A (hydrogen at position R) so that we can not exclude that the density is generated by a mixture of the congeners. Moreover, biochemical experiments using either the purified natural congeners (Brandi et al., 2006b) or chemically synthesised GE81112 congeners A and B1 both show antimicrobial activity, and SAR studies indicate that the NH2 group present at position 2 in AA3 is not essential (Jürjens et al., 2018; Zwick et al., 2021; Schuler et al., 2023). For these reasons and for simplicity, we have modelled the GE81112A congener and discussed it below. In all cryoEM maps from the three independent datasets, GE81112 is observed in the same position, such that a comparison of the binding pocket residues yields RSMD values between 0.924 and 1.138 (297 atom pairs; **Supplemental Figure S4**). This indicates that the presence of tRNA and initiation factors has little effect on the conformation of the binding pocket. In the high-resolution Complex 5 structure, we do notice some evidence for an alternative conformation in U793 in h24 (**Supplemental Figure S5D)**, which is not seen in complex 1-4. This could be due to the lower map resolution in Complex 1-4 or related to the model arising from a 70S ribosome (Complex 5) rather than a 30S initiation complex (Complex 1-4). We note this, however, as chemical probing (Dallas and Noller, 2001) indicates a connection between IF3 and U793, and this alternative conformation could be important for stalling IF3 in position 1 in the presence of GE81112, if the lack of the alternative conformation is related to the lower resolution of complexes 1-4.

**Figure 2:**
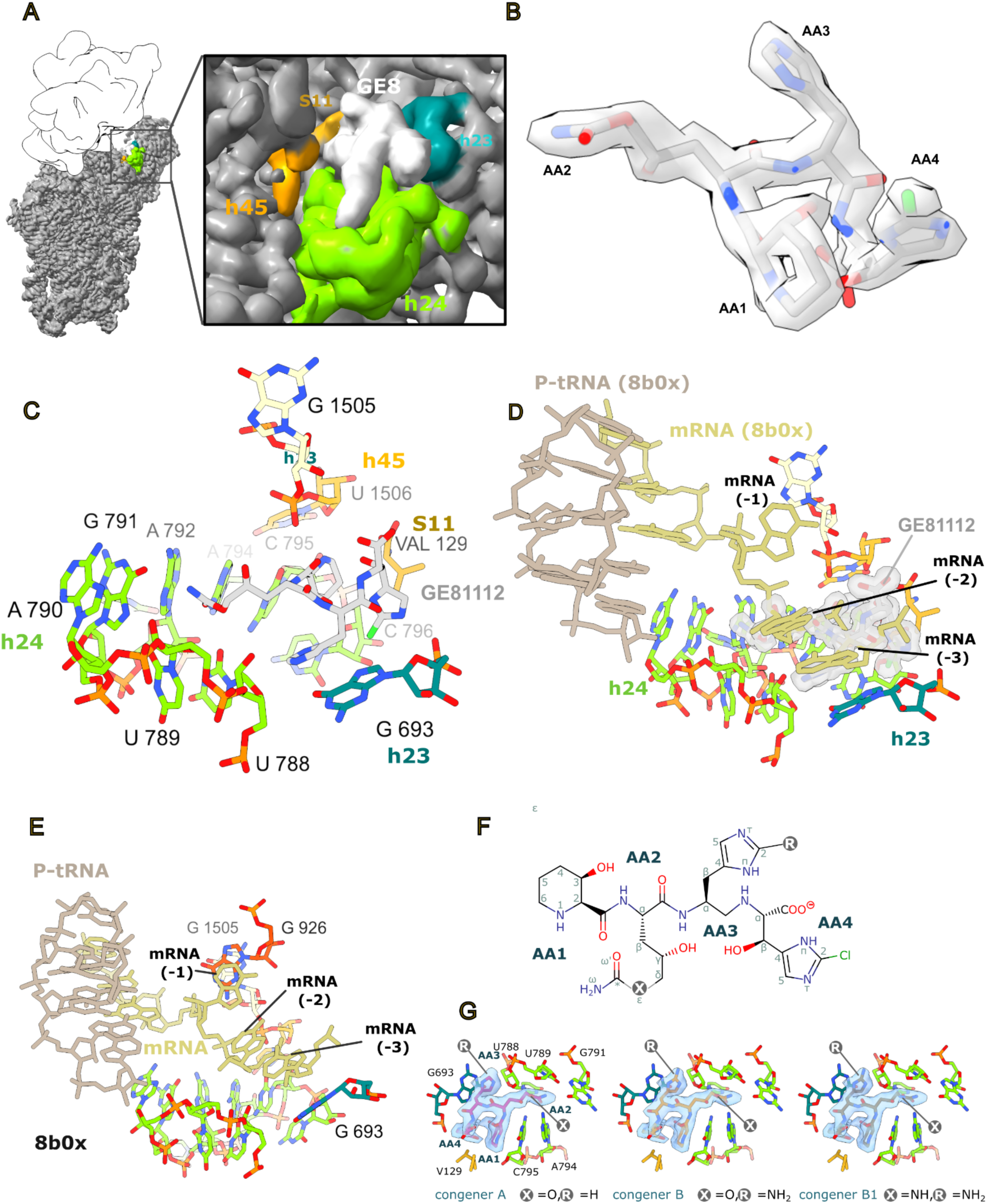
GE81112 binding Site. (**A**) Overview of GE81112 binding site on 30S subunit (Complex 5). (**B**) GE81112 (congener GE81112A) fit to the sharpened complex 5 (cryosparc sharpening) cryoEM map. (**C**) The GE81112 binding pocket overlaps with the mRNA channel of the E-site and is formed by 16S rRNA h24, h23, and h45 and the ribosomal protein (r-protein) S11. (**D**) Aligning the GE81112 model (complex 5, GE81112 is rendered as surface) with a PDB model for a 70S ribosome bound by P-tRNA and mRNA (PDB 8b0x, (Fromm et al., 2023)) shows a clash between GE81112 and the mRNA upstream of the start codon (position -2 and -3 at the mRNA E-site). (**E**) The E-site mRNA is shown on the 8b0x structure in the absence of GE81112 illustrating how the mRNA residues are positioned relative to 16S rRNA residues. For example, in this structure, the -1 residue is stacked on G926, and the -3 residue is positioned over G693. (**F**) 2D chemical structure of GE81112 with the positions that are different in the three congeners marked with an R and an X. The identity of R and X are given in panel **F**. (**G**) The three congeners (labelled in the figure panels) are shown inside the cryoEM density (complex 5; unsharpened).

As noted above (**Figure 2C**), the GE81112 binding site involves only the rRNA (h23, h24, and h45) and r-protein S11 and does not involve any residues from the initiation factors, as seen in complexes 1 and 2. Similarly, it does not contact the fmet-tRNA. Note that in our cryoEM maps (Complexes 1-4), only mRNA residues involved in base-pairing with the tRNA in the P-site show strong density. In the lower-resolution Complex 1 map, there is density with a low local resolution that suggests the mRNA (-1 position) approaches GE81112 such that the mRNA backbone (-1 position) is between G926 and GE81112 (**Supplemental Figure S5C**). G926 is a universally conserved residue that is part of hinge 1, a motif involved in 30S head movement (Mohan et al., 2014). To better understand if GE81112 could potentially interfere with the mRNA outside of the P-site, we aligned a model of a 70S ribosome bound by a P-tRNA and mRNA (PDB 8b0x; (Fromm et al., 2023)) to the complex 5 model. As seen in **Figures 2D** and **E**, residues in the mRNA upstream of the start codon, -2 and -3 (E-site), do clash with GE81112, suggesting that GE81112 interferes with the placement of the mRNA. Interfering with the placement of upstream mRNA residues -2 and -3 (as seen in complex 1-4), agrees with a previous study showing that GE81112 has only a marginal effect on the binding kinetics of model mRNAs but does affect hydroxyl radical cleavage of the mRNA, particularly upstream of the start codon (Fabbretti et al., 2016). We propose that GE81112 prevents proper mRNA accommodation by binding in the E-site portion of the mRNA channel, specifically, we observe that AA3 of GE81112 stacks on G693, a position that is generally occupied by the -3 mRNA residue (Hoffer et al.)(**Figure 2D** and **E**). As seen in **Supplemental Figure S5C**, the cryoEM map suggests the backbone of the -1 residue is positioned near G926. The importance of G926 to the mechanism of action of GE81112 is supported by the previous observation that mutations in G926 reduce the sensitivity of *in vitro* mRNA translation experiments to GE81112 (Maio et al., 2016).

### Interaction of GE81112 with the 30S subunit

Consistent with the hydrophilic nature of GE81112, it makes extensive hydrogen bonds with the 30S subunit. As seen in **Figure 3**, AA1-4 form an extensive hydrogen bond network (shown in yellow) with 16S rRNA residues in helices: h23 (residue G693), h24 (residues U788, U789, A794, and C795), h45 (residue U1506), and the C-terminal carboxyl moiety of ribosomal protein S11 (residue Val129). Moreover, the interaction of GE81112 with the 30S involves a parallel shifted aromatic n stacking (dashed lines in grey-blue) of the imidazol ring of AA3 with the base of G693, and several CH-n interactions (coloured in red/crimson), including those of the methylen moieties of the 3-hydroxy piperidine ring and the isopropyl part of valine 129 with the base of C795 and the imidazol ring of residue AA1 (**Figure 3)**. Ligand binding is favoured by the entropic gains from the displacement of at least three structural waters as identified in the high-resolution X-ray structure of vacant *E.coli* ribosome (PDB ID 4YBB; (Noeske et al., 2015)). The interaction of GE81112 with h23 and h24 is consistent with the chemical probing results of Brandi et al. (Brandi et al., 2006a), showing that G693 is protected from kethoxal modification while C795, and to a lesser extent, A792, A794, C796 all experience changes in dimethyl sulphate reactivity.

**Figure 3:**
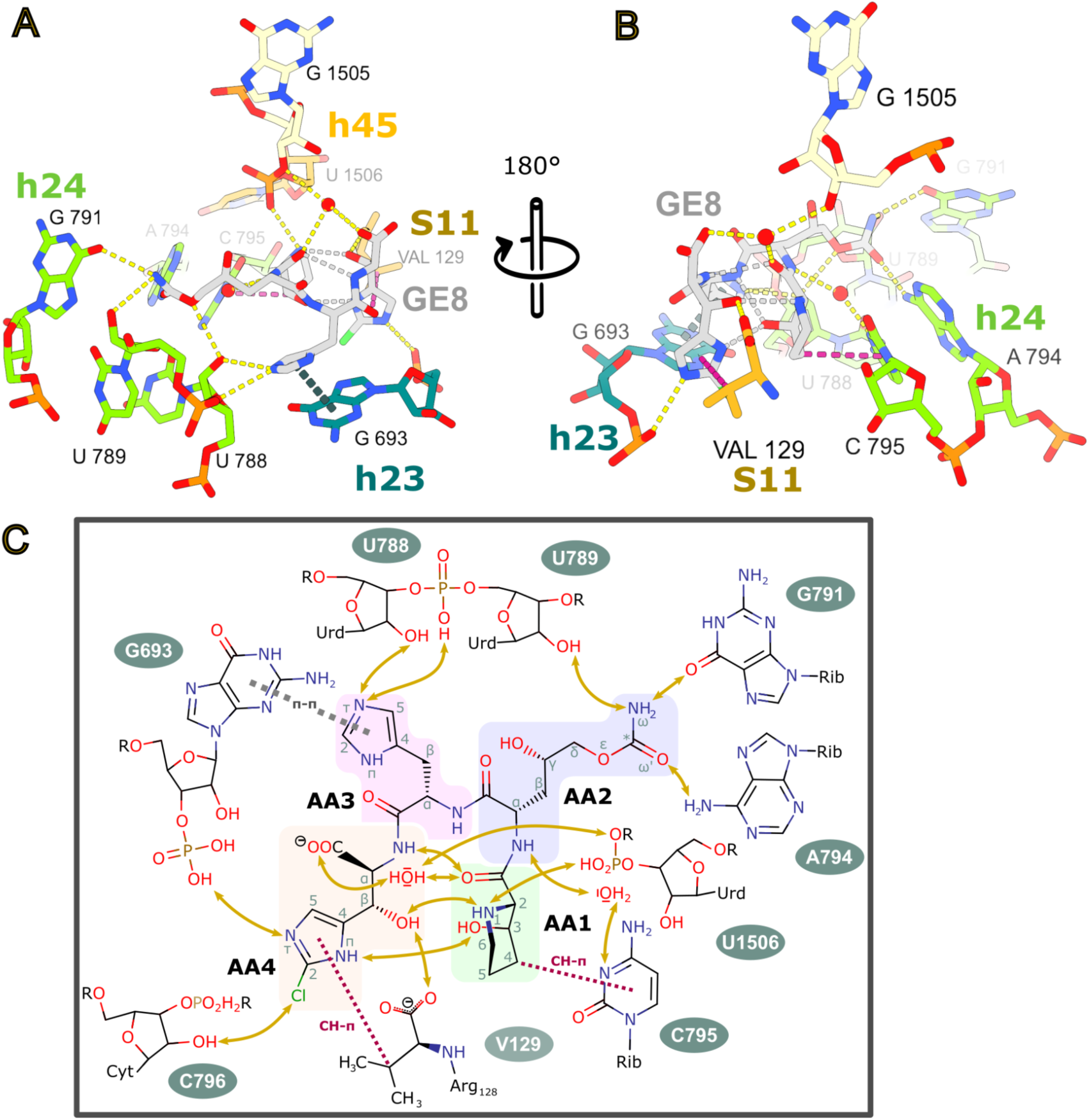
Interaction of GE81112 with the 30S binding pocket. (**A-B)** Two views of the GE81112 (GE8) binding pocket highlighting interactions inferred from geometric constraints. Intermolecular hydrogen bonds are yellow, intramolecular hydrogen bonds are grey, CH-n interactions are crimson, and n stacking interactions are grey-blue. (**C**) The interactions of GE81112 with elements of its binding pocket are summarised in a 2D schematic representation. The amino acid monomers are highlighted: 3-hydroxy-L-pipecolic acid (AA1, green), 4-hydroxy-L-citrulline (AA2, violet), O-carbamoyl-α-amino-dihydroxyvaleric acid, 2-amino-L-histidine (AA3, pink) and β-hydroxy-2-chloro-L-histidine (AA4, cream).

In total, GE81112 is observed to potentially make five direct base-specific interactions (G693, G791, A794, C795 and Val129) and eight direct interactions with sequence-independent groups in the RNA and protein backbone (**Figure 3C)**. The high number of base-specific interactions could afford some degree of targeting GE81112 derivates to specific bacterial strains. However, it should be noted that the rRNA bases involved are highly conserved (G693: 93.52%, G791: 99.88%, A794: 99.69%, C795: 99:86%)^1^. These base-specific interactions also render GE81112 activity sensitive to rRNA mutations. For example, it has been shown that A794G and A794U mutations alter the *in vitro* activity (e.g. mRNA translation) of GE81112 (Maio et al., 2016). As seen in **Figure 2**, these mutations would alter the interaction with AA2, as the positioning of a keto moiety (instead of the 6-amino group of A794) close to the carbamoyloxy- (or carbamoylamino in the case of congener B1) moiety of AA2 would cause a repulsive interaction.

The backbone conformation of the tetrapeptide ligand in the bound state closely resembles a beta-turn of type I (Rose et al., 1985), with various polar moieties of residue AA1 and AA4 stabilising this motif. There are four direct or water-mediated hydrogen bonds between AA1 and AA4, with several involving the 3-hydroxy group of the pipecolic acid (residue AA1) and the beta hydroxy moiety of the chlorinated beta-hydroxyhistidine (residue AA4). This is significant as the SAR studies of Zwick *et al*. (Zwick et al., 2021) demonstrated the “paramount importance” of these groups for antimicrobial activity. If these hydrogen bonds maintain the conformation of GE81112 in solution, they would lower the entropic costs for binding to the 30S subunit, potentially explaining their importance for antimicrobial activity.

### GE81112 shares an overlapping binding site with other initiation-targeting antibiotics

Like GE81112, there are several other chemically unrelated antibiotics, amicoumacin (Polikanov et al., 2014a), edeine (Pioletti et al., 2001), emetine (Wong et al., 2014), kasugamycin (Schluenzen et al., 2006; Schuwirth et al., 2006), and pactamycin (Brodersen et al., 2000; Polikanov et al., 2014a) that bind within the mRNA channel. In **Figure 4**, the structures of these drugs have been aligned with the GE81112 model to highlight the proximity of their binding site. Specifically, there is a substantial overlap between the binding site of GE81112 and amicoumacin (**Figure 4B**), emetine (**Figure 4D**) and pactamycin (**Figure 4G-H**), while edeine and kasugamycin (**Figure 4C, E, F**) bind more outside the GE81112 pocket. Amicoumacin, for example, binds to this site but stabilises the mRNA, affecting both translation and initiation (increased formation of erroneous 30S ICs)(Polikanov et al., 2014a; Maksimova et al., 2021). These drugs also impact mRNA function in a variety of ways. Edeine is a peptide antibiotic that blocks 30S formation by preventing P-tRNA binding (Dinos et al., 2004). Emetine is an anti-protozoan drug affecting mRNA/tRNA translocation (Wong et al., 2014). Collectively, this establishes that the h24/h23 binding site, near the E-site mRNA codon, is a hotspot for disturbing mRNA function as it affects binding and recognition of codon and anticodon in both the P and A sites.

**Figure 4:**
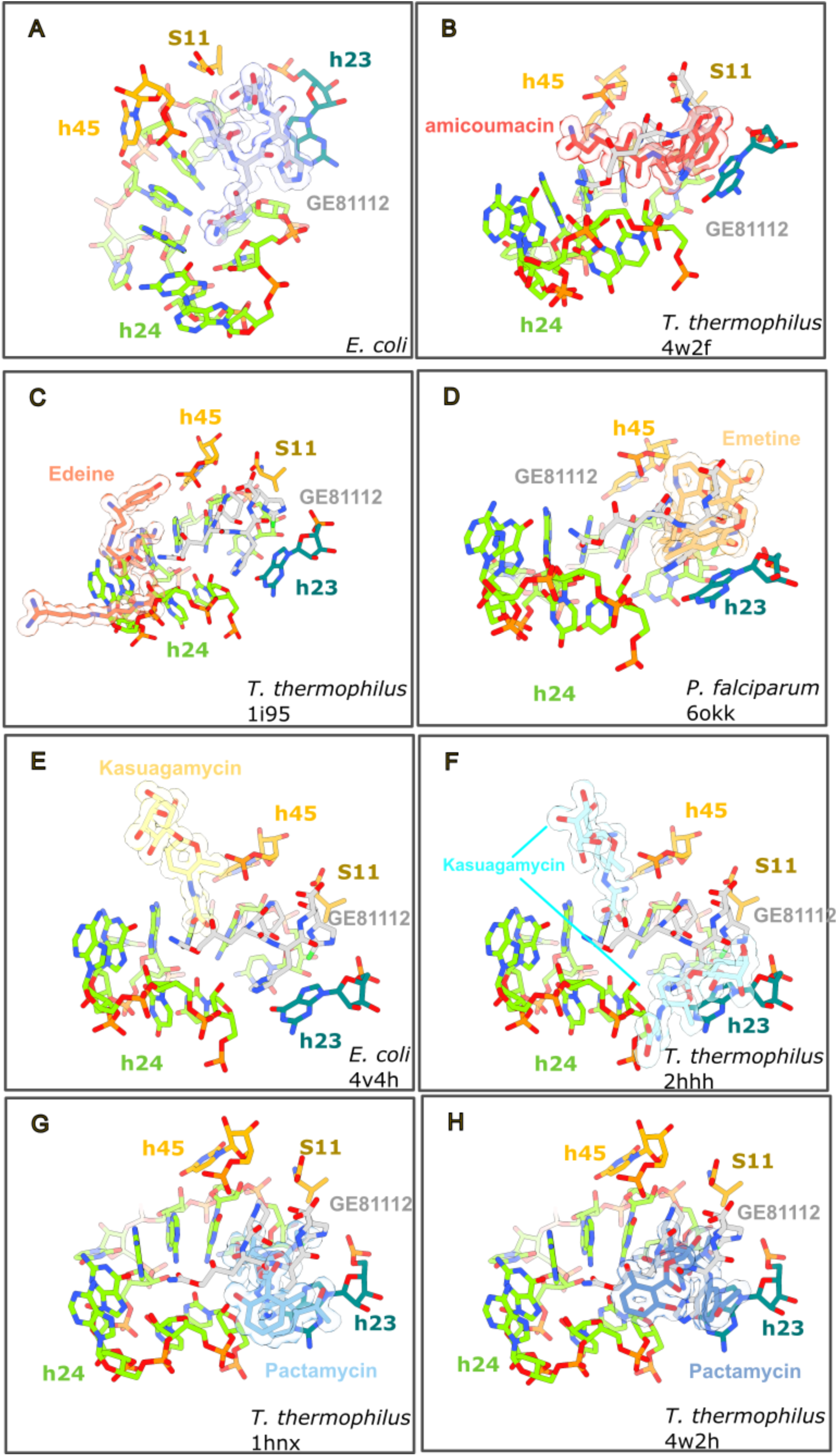
The GE81112 binding pocket is shared by other mRNA-targeting antibiotics. PDB models (**B**) 4w2f, (**C**) 1i95, (**D**) 6okk, (**E**) 4v4h, (**F**) 2hhh, (**G**) 1hnx and (**H**) 4w2h have been aligned to the GE81112 binding pocket (**A**) using residues within 15 Å of GE81112. The orientation in each panel was selected to highlight the relative positions of GE81112 and the other antibiotics.

## DISCUSSION

In this study, we present five structures from 3 independent samples that describe the interaction between GE81112 and the *E. coli* ribosome in the context of the initiation of protein synthesis. These structures show GE81112 binds to the 30S subunit within the mRNA channel in a pocket, termed hereafter ecGE81112 binding site, formed by helices h23, h24, and h45 of 16S rRNA, as well as ribosomal protein S11 (**Figure 2**). This is distinct from our previously identified site in the crystal structure of *T. thermophilus* 30S subunit (ttGE81112), which is in proximity of the distorted h6 of a symmetry-related 30S subunit, which is proposed to mimic the anticodon stem-loop of a P-tRNA (Fabbretti et al., 2016). In the crystal, despite most of the elements that constitute the ecGE8112 binding site being present, no density was observed at this position for GE81112. In this respect, we notice that one difference between the two binding sites is represented by the C-terminal tail of S11. It is positioned differently in the *T. thermophilus* crystal, possibly because *T. thermophilus* lacks S21. Moreover, the C-terminal valine residue (V129), which is conserved among various bacterial species, is absent in *T. thermophilus* (**Supplemental Figure S6A**). Consequently, S11 may not contribute to GE81112 binding in *T. thermophilus* by forming CH-n interactions with GE81112 via V129, as seen in *E. coli* (**Figure 3**). Regarding the ttGE81112 binding site, it is not formed in the *E. coli* cryoEM maps obtained in this study. This may be due to the constraints imposed for the formation of the crystals, which include limiting the freedom of the ASL mimic. Also, in the crystal structure, to understand the basis of the altered h6 (ASL mimic) conformation observed, even in the Fo-Fc map, we employed a bulk solvent modelling protection approach (Polikanov et al., 2014b; Liebschner et al., 2017). In the resulting map, we could account for only part of the density generated by ordering of the C-terminal tail of S13 (K121-K126), which is usually disordered in the crystal structures. The remaining part of the density could be explained by the presence of GE81112 and, as indicated in our previous work, “although the ligand GE81112 and S13 can be putatively modelled into the electron density, it is not possible at the resolution of the map to distinguish the last C-terminal residues of S13 unambiguously from the four nonproteinogenic amino acid moieties of GE81112, and therefore alternative arrangements are possible” (Fabbretti et al., 2016). In this respect, one possibility not considered earlier would have been to model S13 in multiple conformations although this flexibility would be at odds with the stable conformation of the ASL mimic observed even in the Fo-fc map (Fabbretti et al., 2016). Furthermore, considering the C-terminal residues of S13 (K121-K126) as part of the ttGE81112 binding pocket, it is relevant to note that S13 in *E. coli* is shorter, composed of just 118 amino acids (**Supplemental Figure S6B**), and this may also explain why this binding pocket is not observed in the *E. coli* system. Finally, considering the system explored in this study is less artificial, the resolution obtained in the cryoEM data is much higher, the ecGE81112 binding site (agrees best with the chemical probing data (Brandi et al., 2006a; Fabbretti et al., 2016) and the sensitivity of GE81112 *in vitro* activity to rRNA mutations (Maio et al., 2016), we consider it consistent with being the biologically active site.

As seen in **Figure 2**, the map presented here nicely accommodates the chemical scaffold of GE81112, and modelling indicates the drug forms an extensive hydrogen bond network with the 16S rRNA and r-protein S11, while its cyclic moieties engage in n-stacking and CH-n interactions, particularly with G693 and C795 (**Figure 3**). Notably, the cryo-EM analysis reveals that the initiation factors and tRNA do not directly interact with GE81112, suggesting its inhibitory mechanism stems primarily from GE81112 being positioned in the mRNA binding channel. In this position, it prevents the mRNA, the -2 and -3 positions, from being accommodated in the groove of h24; specifically, AA3 in GE81112 replaces the -3 mRNA residue in stacking on G693 (Hoffer et al.) (**Figure 2D** and **E**). The disruption of the upstream mRNA path (-2 and -3 residues, Complex 1-4, **Supplemental Figure S7**) agrees with previous hydroxyl radical probing experiments that show GE81112 alters the interaction between the upstream residues of the mRNA and 30S subunit (Fabbretti et al., 2016). It should be noted that the solvent-exposed face of GE81112 contains carbonyl moieties that would have a repulsive effect on the backbone of the mRNA, contributing to the -2 and -3 residues being disordered in our maps. The -1 mRNA residue is only partially disordered in our Complex C map; there appears to be density for the backbone but not the nucleobase (**Supplemental Figure S1E** and **S5C**). Importantly, the density suggests the -1 residue of the mRNA is positioned near G926 (**Supplemental Figure S1E a**nd **S5C**), a universally conserved bulged base in the h28 (the neck) that Mohan *et al*. propose is the point (hinge 1) that the head pivots around during translocation (Mohan et al., 2014). The nodding/swivelling movement of the head has been shown by Lopez *et al*. and Hussain *et al*. (Hussain et al., 2016; López-Alonso et al., 2017) to alter the configuration of the P-site and reposition the codon-anticodon complex during the conversion of early preIC complex to 30S IC complexes, where the tRNA moves from an unaccommodated to an accommodated state. We hypothesise that GE81112 could disfavour correct head rotation by influencing hinge 1 via G926. There are two routes for GE81112 to influence G926. The first would be via the -1 mRNA residue; namely, in the complex C structure, the E-site portion of the mRNA is largely disordered and does not make defined interactions, for example, stacking with G926 like seen in other structures (Nishima et al., 2022; Hoffer et al.). The second route could exploit the interaction of GE81112 with the phosphate connecting G1505 and U1506 (**Figure 3**), because G1505, in fact, stacks on G926 from the other face. A role of G926 in GE81112 activity is suggested by the work of Maio *et al*. where mutations in G926 reduce the sensitivity of *in vitro* mRNA translation experiments to GE81112 (Maio et al., 2016).

One interesting aspect of the GE81112 model is the close proximity of AA1 and AA4, which is maintained by an extensive network of intramolecular hydrogen bonds (**Figure 3B**). This network includes the 3-hydroxy group of the pipecolic acid (residue AA1) and the β-hydroxy moiety of the chlorinated β-hydroxyhistidine (residue AA4), which Zwick *et al*. (Zwick et al., 2021) demonstrated to be critical for the antimicrobial activity. This close association of AA1 and AA4 could facilitate cyclising the GE81112 scaffold. This is significant because cyclic peptide antibiotics are playing an increasing role in modern drug discovery efforts (Lai et al., 2022). Cyclic peptides can increase metabolic stability due to reduced degradation, while their reduced conformational flexibility can contribute entropically to increase target binding (Lai et al., 2022). A cyclic GE81112 derivative could also alter the uptake through the OPP system, potentially bypassing a significant pathway for GE81112 resistance (Maio et al., 2016; Schuler et al., 2023). The structural insights from this study provide a foundation for optimising GE81112 derivatives and exploring their potential in combating bacterial infections.

## MATERIAL AND METHODS

### Assembly of the pre*IC*

Complexes for both datasets were prepared independently under identical conditions. *Escherichia coli* ribosomes, ribosomal subunits, and translational factors were prepared as previously described (López-Alonso et al., 2017). Using purified components, the 30SIC was assembled by incubating the 30S ribosomal subunits with GE81112 for 10 min at 37°C in Buffer IC (10 mM Tris-HCl (pH 7.7), 7 mM MgCl_2_, and 60 mM NH_4_CH_3_COO) prior to addition of the mRNA construct containing a model Shine-Dalgarno sequence (AAG UUA ACA GGU AUA CAU ACU AUG UUU ACG AUU ACU ACG AUC), fMet-tRNA, GTP and IF1, IF2, IF3, and continuing the incubation at 37°C for an additional 10 min. Throughout this incubation, buffer conditions are kept constant, and the final factor concentrations are 1 µM 30S, 20 µM GE81112, 4 µM mRNA, 2.8 µM fMet-tRNA, 50 µM GTP, 3.2 µM IF1, 1.6 µM IF2, and 3.2 µM IF3.

### Vitrification and Electron microscopy of preIC complex

The 30SIC sample was vitrified using a Vitrobot (Thermo Fisher Scientific, TFS) by diluting the reconstituted 30S IC in Buffer IC containing 20 µM GE81112, 4 µM mRNA, 2.8 µM fMet-tRNA and 50 µM GTP onto Quantifiol R2/2 grids. Automated data acquisition (EPU software, TFS) was performed at NeCEN (Netherlands Centre for Electron Nanoscopy; dataset 1) and eBIC (Diamond Light Source, UK; dataset 2) with a Titan Krios microscope (FEI) at 300 kV equipped with a direct detector (**Supplemental Table 1**). For Dataset 1, 3745 movies were collected at a pixel size of 1.086 A and defocus ranging from 0.4 µm to 3.8 µM. For Dataset 2, 6172 movies were collected, each containing 19 frames at a pixel size of 1.113 A. More detailed imaging conditions are presented in **Supplemental Table 1**.

### Vitrification and Electron microscopy of 70S-GE81112 complex

The GE81112-70S complex we prepare by co-incubating (10 min, ice) 100 nM 70S ribosomes with 20 µM GE81112 (AdipoGen Life Science) in buffer consisting of 10 mM Tris pH 8.0 14 mM MgAc and 60 mM KCl. The sample was vitrified using a Vitrobot (TFS) on Quantifoil R1.2/1.3 (300 mesh) grids. Automated data acquisition (EPU software, TFS) was performed at the Basque Resource for Electron Microscopy (BREM) on a 300 kV X-FEG Krios G4 transmission electron cryo-microscope (TFS) and a Gatan K3 direct detector (**Supplemental Table 1**). For Dataset 3, 19,615 movies were collected, each fractionated into 40 frames at a pixel size of 0.8238 Å/px and a total exposure dose of 49.3 e-/A^2^.

### Single Particle Analysis

#### Dataset 1

Motion correction was performed within RELION 3.1 (Zivanov et al., 2020)using the dose weighting and patch (5 x 5) options. Contrast transfer function (CTF) estimation for each aligned micrograph was performed using CTFFind (Rohou and Grigorieff, 2015). 232,679 projection images of 30S particles were picked using crYOLO (Wagner et al., 2019). Initially, particles were rescaled and extracted with a pixel size of 3.01 Å, and the dataset was cleaned by using RELION 2D Classification (100 classes). Subsequently, the well-aligning particle projections (total of 193743) were used to generate an initial model. The projections were then refined in several cycles (using the initial mode to start), yielding a 5.2 Å map where the body region was visually more defined than the head region. This prompted us to utilise a multi-body refinement (head and body mask), which improved the resolution of the body region to 4.7 Å. The cryoEM map of the 30S body region was then used to initiate CTF refinement (beam tilt, anisotropic magnification, per particle defocus fitting, per micrograph astigmatism) and Bayesian polishing (Zivanov et al., 2022). CTF correction was applied to the polished particles to reduce their window size and speed up processing. After polishing, the map was refined to 3.7 Å. 3D classification was performed to remove poorly aligning particles, such that we finally retained a subset of 46163 that yielded the final 3.8 Å map. Further, multibody refinements or 3D classifications did not improve the map as judged visually or by resolution estimates. The FSC (Fourier shell correlation) plots for the consensus refinement, the final map and local resolution estimates are shown in **Supplemental Figure 1**.

#### Dataset 2

All data processing steps were performed within RELION 3.0 or 3.1 (Zivanov et al., 2018, 2020). 6172 movies (**Supplemental Figure 2A**) were imported and motion-corrected with Relion. Defocus estimation was performed with Gctf v1.06 (Zhang, 2016), and subsequently, micrographs with defocus values between -0.5 and -3.25 µm were selected. Particles were picked with cryolo ((Wagner et al., 2019); 662792 particles) and initially extracted at a pixel size of 2.226 Å/px. These particles were cleaned by 2D classification (100 classes; **Supplemental Figure 2A**), such that 397515 particles were retained and used to generate an initial model. Further cleaning was performed with 3D classification so that 304619 particles were kept and finally refined with a pixel size of 1.113 Å/px to generate a 3.2 Å cryoEM map. The cryoEM map of the 30S body region was then used to initiate CTF refinement and Bayesian polishing (Zivanov et al., 2022). Subsequent refinement led to a 2.95 Å cryo map. 3D classification (4 classes, no image alignment, T=16) was performed on these particles using a mask centred on the density of IF1, IF3 and the tRNA anticodon arm. This yielded three well-defined volumes that were subsequently refined to 3.3 Å (Complex 2), 3.3 Å ( Complex 3) and 3.1 Å (Complex 4). Finally, a multibody refinement was performed where body 1 corresponded to the 30S body and body 2 to the 30S head region. When present in the map, the best multibody refinements were obtained when the tRNA was included in the body 2 mask and the IFs in the body 1 mask. The FSC plots for the multibody maps, the final maps and local resolution estimates are shown in **Supplemental Figure 1**.

#### Dataset 3

All image processing steps were performed within the CryoSPARC and CryoSPARC Live software package (Punjani et al., 2017). Movies were imported to CryoSPARC, patch motion corrected (**Supplemental Figure S3A**), and the contrast transfer function was estimated. The micrographs were curated based on estimated resolution, defocus values, and motion parameters. The particles were windowed out and downsampled to a box size of 128 pixels x 128 pixels for initial cleaning using 2D classification (**Supplemental Figure S3B**). The best classes resembling 70S, 50S, and 30S ribosomes were selected, finally corresponding to 1,816,931 particles. These particles were used in an *ab initio* reconstruction job (six classes). Subsequently, heterogeneous refinement and 3D classification jobs (Punjani and Fleet, 2021) were used to separate the input projections into poorly aligning, 30S (29,389), 50S (190,467) and 70S (1,331,197) classes. The 70S particles were re-extracted at 512 pixels x 512 pixels and refined, yielding a cryoEM map at 2.05Å resolution (70S). Beam tilt correction (Zivanov et al., 2020) and reference-based motion correction were performed, and after a subsequent non-uniform refinement, using a 50S mask, a 70S cryoEM map at 1.85Å resolution was obtained (1,329,758 particles; under a 50S mask; **Supplemental Figure S3C-D**). A particle subtraction job was performed on these 50S aligned projections to generate projections containing signal from only the 30S body region of the 70S ribosome. After a non-uniform refinement, a cryoEM map of the 30S body region at 2.09Å was obtained (**Supplemental Figure S3E-F,** 1,329,758 particles). Local resolution was estimated with CryoSPARC using the non-uniform refinement half maps (Punjani et al., 2020).

### Cryo-EM model building

The absolute configuration of GE81112 for describing the ligand topology was derived from Jürjens *et al*. (Jürjens et al., 2018). Starting ligand coordinates for refinement were generated using the RDkit package for cheminformatics (RDKit: Open-source cheminformatics. https://www.rdkit.org), and restraint information about its geometry was derived using eLBOW (Moriarty et al., 2009). The PBD structure 4YBB (Noeske et al., 2015) was taken as an initial template for refinement and model building. Individual maps for the head (encompassing rRNA residues C931 to G1386 and the ribosomal proteins S3, S7, S9, S10, S13, S14, and S19) and body domain (residues A1 to C930 and G1387 to A1542 of the 16S RNA, and S3, S7, S9, S10, S13, S14, and S19 r-proteins) were derived from cryo-EM maps obtained by the RELION multi-body refinement (low-pass filtered to the global resolution). After a preliminary rigid body refinement followed by several cycles of manual model-building using Coot (Emsley et al., 2010) and real-space refinement in Phenix (Liebschner et al., 2019)(with secondary structure restraints and Ramachandran restraints) the non-ribosomal components of the complex were added at the final refinement whose starting coordinates were taken either from the PDB (PDBID 6UGG for tRNA) or from structure predictions employing AlphaFold2 (for the initiation factors)(Jumper et al., 2021). Structural fragments of the ribosomal complexes not accounted for by the cryo-EM density, due to their absence or local disorder, were omitted from the final models. The quality of the obtained models was assessed using MolProbity as a part of the Phenix validation tools (Liebschner et al., 2019), and the guanidino carboxy denotation issues were resolved by an in-house script. For figure preparation, ChimeraX (Meng et al., 2023) and Inkscape 1.3 were used.

## ACCESSION NUMBERS

The raw electron microscopy movies, cryo-EM maps, and resulting atomic models are available at the EMPIAR, EMDB, and Protein Data Bank (PDB) under accession codes XXXXX, YYYYY, and ZZZ. Data are available from the corresponding authors upon reasonable request.

## SUPPLEMENTARY DATA

Supplementary Data are available online.

## ACKNOWLEDGEMENT

This work was supported by grant PID2021-122705OB-I00 (to S.R.C and P.F.) funded by MCIN/AEI/10.13039/501100011033 and, as appropriate, by “ERDF A way of making Europe”, by the “European Union” or by the “European Union NextGenerationEU/PRTR”. Support was also provided by the Basque Government grant IT1578-22 (to P.F and R.Z); the MINECO grant CTQ201782222-R (to P.F and S.R.C); the One Health Observatory Lighthouse (HOBE) in Plentzia Bay (TED2021-132109B-C21) by MINECO and the European Union (P.F.). RZ and RP are research fellows supported by the cotutelle program PIF(PAU-UPV/EHU). The authors acknowledge the support and the use of resources of Instruct, a Landmark ESFRI project (specifically Instruct Access Project PID: 1604-73), including the Netherlands Centre for Electron Nanoscopy (NeCEN, Leiden). We acknowledge Diamond for access and support of the Cryo-EM facilities at the UK national electron bio-imaging centre (eBIC), proposal EM17171-2 and bi31586-33, funded by the Wellcome Trust, MRC and BBSRC. Some of this work was performed at the Basque Resource for Electron Microscopy located at Instituto Biofisika (UPV/EHU, CSIC), supported by the Department of Education and the Innovation Fund of the Basque Government, with additional support from MCIN (Recovery, Transformation and Resilience Plan) and the Basque Government “Biotechnology Complementary Plan Applied to Health” with funding from European Union NextGenerationEU (PRTR-C17.I1; PRTR-C17.I01.P01.S13) (AAAA_ACG_AY_2539/22_05). Molecular graphics and analyses performed with UCSF ChimeraX, developed by the Resource for Biocomputing, Visualization, and Informatics at the University of California, San Francisco, with support from National Institutes of Health R01-GM129325 and the Office of Cyber Infrastructure and Computational Biology, National Institute of Allergy and Infectious Diseases.

## Footnotes

1 https://crw2-comparative-rna-web.org/nucleotide-frequency/16s-rrna-model-single-base-frequency/

