## Supplemental Material for "A binding site for the antibiotic GE81112 in the ribosomal mRNA channel"

* Former address in the early stage of the project: Center for Cooperative Research in Biosciences (CIC bioGUNE).

** To whom correspondence should be addressed.

### TABLE OF CONTENTS

#### 1. Supplementary Tables

Supplemental Table S1: EM Data Collection and Image Processing

Supplemental Table S2: Model Validation Statistics for State 1

Supplemental Table S3: Model Validation Statistics for State 2

Supplemental Table S4: Model Validation Statistics for State 3

Supplemental Table S5: Model Validation Statistics for State 4

Supplemental Table S6: Model Validation Statistics for State 5

#### 2. Supplementary Figures

Supplemental Figure S1: Cryo-EM Processing Results for Dataset 1.

Supplemental Figure S2: Cryo-EM Processing Results for Dataset 2.

Supplemental Figure S3: Cryo-EM Processing Results for Dataset 3.

Supplemental Figure S4: Comparison of GE81112 pocket in Complexes 1-5.

Supplemental Figure S5: The initiation complex 1 is in a preIC state.

Supplemental Figure S6: Conservation of r-proteins S11 and S13.

Supplemental Figure S7: Key mRNA, tRNA and 30S interactions are altered or disrupted in the GE81112 pre-initiation complex.

#### SUPPLEMENTARY TABLES

**Supplemental Table S1: EM Data Collection and Image Processing.**

| **Data Collection** | | | |
|---|---|---|---|
| | **EMD-<br>EMPIAR-** | **EMD-<br>EMPIAR-** | **EMD-<br>EMPIAR-** |
| Sample | ***Dataset 2<br>30SpreIC-GE8*** | ***Dataset 1<br>30SpreIC-GE8*** | ***Dataset 3<br>70S-GE8*** |
| Facility | eBIC | NeCEN | BREM |
| Microscope | Titan Krios | Titan Krios | Titan Krios |
| Camera | | K2 | K3 |
| Data Collection Software | EPU (TFS) | EPU (TFS) | EPU (TFS) |
| Voltage (kV) | 300 | 300 | 300 |
| Calibrated Pixel Size (Å) | 1.113 | 1.086 | 0.8238 |
| Total Exposure ($e^{-}$/ Å $^{2}$) | 44.08 | 51 | 49.3 |
| Number of Frames | 19 | 32 | 40 |
| Defocus Range (µm) | -0.5 to -3.25 | -0.4 to -3.8 | -0.1 to -2.5 |
| **Image Processing** | | | |
| Motion Correction Software | motioncorr | motioncorr | cryoSPARC |
| CTF estimation software | CtfFind | CtfFind | cryoSPARC |
| Particle Section | crYOLO | crYOLO | cryoSPARC |
| Micrographs Collected | 6175 | 3745 | 19615 |
| Particles Selected | 451301 | 232679 | 1694913 |
| Classification and Refinement Software | Relion | Relion | cryoSPARC |
| **Model Building** | | | |
| Visualization Software | ChimeraX | ChimeraX | ChimeraX |
| Refinement Software | Phenix, Isolde | Phenix, Isolde | Phenix, Isolde |

**Supplemental Table S2.** Model Validation Statistics for ***State 1***.

| Complex | state 1 (30S) |
|---|---|
| **PDB** | |
| **EMDB** | |
| **Model composition** | |
| Chains | 30 |
| Non-hydrogen atoms | 58837 |
| Protein residues | 3061 |
| Nucleotides | 1616 |
| Ligands | |
| GE81112 | 1 |
| Mg | 150 |
| K | 5 |
| Zn | 1 |
| **RMSD deviations from ideal values** | |
| Bond length (Å) | 0.005 |
| Bond angles (°) | 0.817 |
| **Ramachandran plot (%)** | |
| Favored | 97.35 |
| Allowed | 2.65 |
| Outliers | 0.00 |
| **Other structural quality metrics** | |
| MolProbity score | 1.73 |
| Clash score | 15.33 |
| Rotamer outliers (%) | 0.0 |
| Cb outliers (%) | 0 |
| Cis or twisted non-trans peptide planes (%) | |
| Cis proline | 0 |
| Twisted proline | 0 |
| **ADP (B-factors)** | |
| Iso/Aniso (#) | 58837/0 |
| Min/max/mean | |
| Protein | 8.98/165.7/58.9 |
| Nucleotide | 12.46/191.06/43.93 |
| Ligand | 7.42/140.58/42.7 |

*$Mg^{2+}$ ions, $K^{+}$ ions and water O are present with chain IDs 5 and 6, and 7, respectively, ligand as residue number 1601 in chain A.

**Supplemental Table S3.** Model Validation Statistics for ***State 2***.

| Complex | state 2 (Head) | state 2 (Body) |
|---|---|---|
| **PDB** | | |
| **EMDB** | | |
| **Model composition** | | |
| Chains | 13 | 18* |
| Non-hydrogen atoms | 18982 | 34842 |
| Protein residues | 1102 | 1462 |
| Nucleotides | 478 | 1078 |
| Ligands | | |
| GE81112 | 0 | 1 |
| Mg | 49 | 93 |
| K | 0 | 4 |
| Zn | 1 | 0 |
| **RMSD deviations from ideal values** | | |
| Bond length (Å) | 0.005 | 0.006 |
| Bond angles (°) | 0.765 | 0.891 |
| **Ramachandran plot (%)** | | |
| Favored | 98.00 | 98.61 |
| Allowed | 2.00 | 1.39 |
| Outliers | 0.00 | 0.00 |
| **Other structural quality metrics** | | |
| MolProbity score | 1.55 | 1.31 |
| Clash score | 10.65 | 5.64 |
| Rotamer outliers (%) | 0.00 | 0.00 |
| Cb outliers (%) | 0 | 0 |
| Cis or twisted non-trans peptide planes (%) | | |
| Cis proline | 0 | 0 |
| Twisted proline | 0 | 0 |
| **ADP (B-factors)** | | |
| Iso/Aniso (#) | 18982/0 | 34843/0 |
| Min/max/mean | | |
| Protein | 30.00/157.81/81.65 | 8.98/96.22/32.28 |
| Nucleotide | 20.00/191.06/71.66 | 12.46/84.24/31.02 |

*$Mg^{2+}$ ions, $K^{+}$ ions and water O are present with chain IDs 5 and 6, and 7, respectively, ligand as residue number 1601 in chain A.

**Supplemental Table S4.** Model Validation Statistics for ***State 3***.

| Complex | state 3 (Head) | state 3 (Body) |
|---|---|---|
| **PDB** | | |
| **EMDB** | | |
| **Model composition** | | |
| Chains | 13 | 15* |
| Non-hydrogen atoms | 18869 | 33359 |
| Protein residues | 1091 | 1280 |
| Nucleotides | 478 | 1078 |
| Ligands | | |
| GE81112 | 0 | 1 |
| Mg | 44 | 92 |
| K | 0 | 6 |
| Zn | 1 | 0 |
| **RMSD deviations from ideal values** | | |
| Bond length (Å) | 0.007 | 0.006 |
| Bond angles (°) | 0.942 | 0.907 |
| **Ramachandran plot (%)** | | |
| Favored | 96.00 | 98.73 |
| Allowed | 4.00 | 1.27 |
| Outliers | 0.00 | 0.00 |
| **Other structural quality metrics** | | |
| MolProbity score | 2.21 | 1.32 |
| Clash score | 28.83 | 5.84 |
| Rotamer outliers (%) | 0.43 | 0.10 |
| Cb outliers (%) | 0 | 0 |
| Cis or twisted non-trans peptide planes (%) | | |
| Cis proline | 0 | 0 |
| Twisted proline | 0 | 0 |
| **ADP (B-factors)** | | |
| Iso/Aniso (#) | 19039/0 | 33357/0 |
| Min/max/mean | | |
| Protein | 45.15/165.07/82.26 | 8.98/44.07/24.24 |
| Nucleotide | 20.00/191.06/71.47 | 12.46/84.24/31.02 |

*$Mg^{2+}$ ions, $K^{+}$ ions and water O are present with chain IDs 5 and 6, and 7, respectively, ligand as residue number 1601 in chain A.

**Supplemental Table S5.** Model Validation Statistics for ***State 4***.

| Complex | state 4 (Head) | state 4 (Body) |
|---|---|---|
| **PDB** | | |
| **EMDB** | | |
| **Model composition** | | |
| Chains | 11 | 15* |
| Non-hydrogen atoms | 18486 | 33344 |
| Protein residues | 1100 | 1278 |
| Nucleotides | 456 | 1078 |
| Water | 0 | 0 |
| Ligands | | |
| GE81112 | 0 | 1 |
| Mg | 42 | 95 |
| K | 0 | 12 |
| Zn | 1 | 0 |
| **RMSD deviations from ideal values** | | |
| Bond length (Å) | 0.005 | 0.005 |
| Bond angles (°) | 0.826 | 0.575 |
| **Ramachandran plot (%)** | | |
| Favored | 98.52 | 98.3 |
| Allowed | 1.48 | 1.67 |
| Outliers | 0.00 | 0.00 |
| **Other structural quality metrics** | | |
| MolProbity score | 1.46 | 1.49 |
| Clash score | 8.62 | 9.20 |
| Rotamer outliers (%) | 0.00 | 0.00 |
| Cb outliers (%) | 0.00 | 0.00 |
| Cis or twisted non-trans peptide planes (%) | | |
| Cis proline | 0 | 0 |
| Twisted proline | 0 | 0 |
| **ADP (B-factors)** | | |
| Iso/Aniso (#) | 18486/0 | 33344/0 |
| Min/max/mean | | |
| Protein | 45.15/154.95/81.47 | 8.98/42.43/24.10 |
| Nucleotide | 45.08/191.06/72.27 | 12.46/84.24/31.02 |

* Zn and Mg ions are present with chain ID 3, ligand as as residue number 1601 in chain A.

**Supplemental Table S6.** Model Validation Statistics for ***State 5***.

| Complex | state 5 (Body) |
|---|---|
| **PDB** | |
| **EMDB** | |
| **Model composition** | |
| Chains | 15* |
| Non-hydrogen atoms | 33344 |
| Protein residues | 1278 |
| Nucleotides | 1078 |
| Water | 1881 |
| Ligands | |
| GE81112 | 1 |
| Mg | 99 |
| K | 35 |
| Zn | 0 |
| **RMSD deviations from ideal values** | |
| Bond length (Å) | 0.007 |
| Bond angles (°) | 0.834 |
| **Ramachandran plot (%)** | |
| Favored | 97.93 |
| Allowed | 2.07 |
| Outliers | 0.00 |
| **Other structural quality metrics** | |
| MolProbity score | 1.48 |
| Clash score | 8.47 |
| Rotamer outliers (%) | 0.10 |
| Cb outliers (%) | 0.00 |
| Cis or twisted non-trans peptide planes (%) | |
| Cis proline | 0 |
| Twisted proline | 0 |
| **ADP (B-factors)** | |
| Iso/Aniso (#) | 33344/0 |
| Min/max/mean | |
| Protein | 8.98/42.43/24.10 |
| Nucleotide | 12.46/84.24/31.02 |

* Zn and Mg ions are present with chain ID 3, ligand as as residue number 1601 in chain A.

#### SUPPLEMENTARY FIGURES

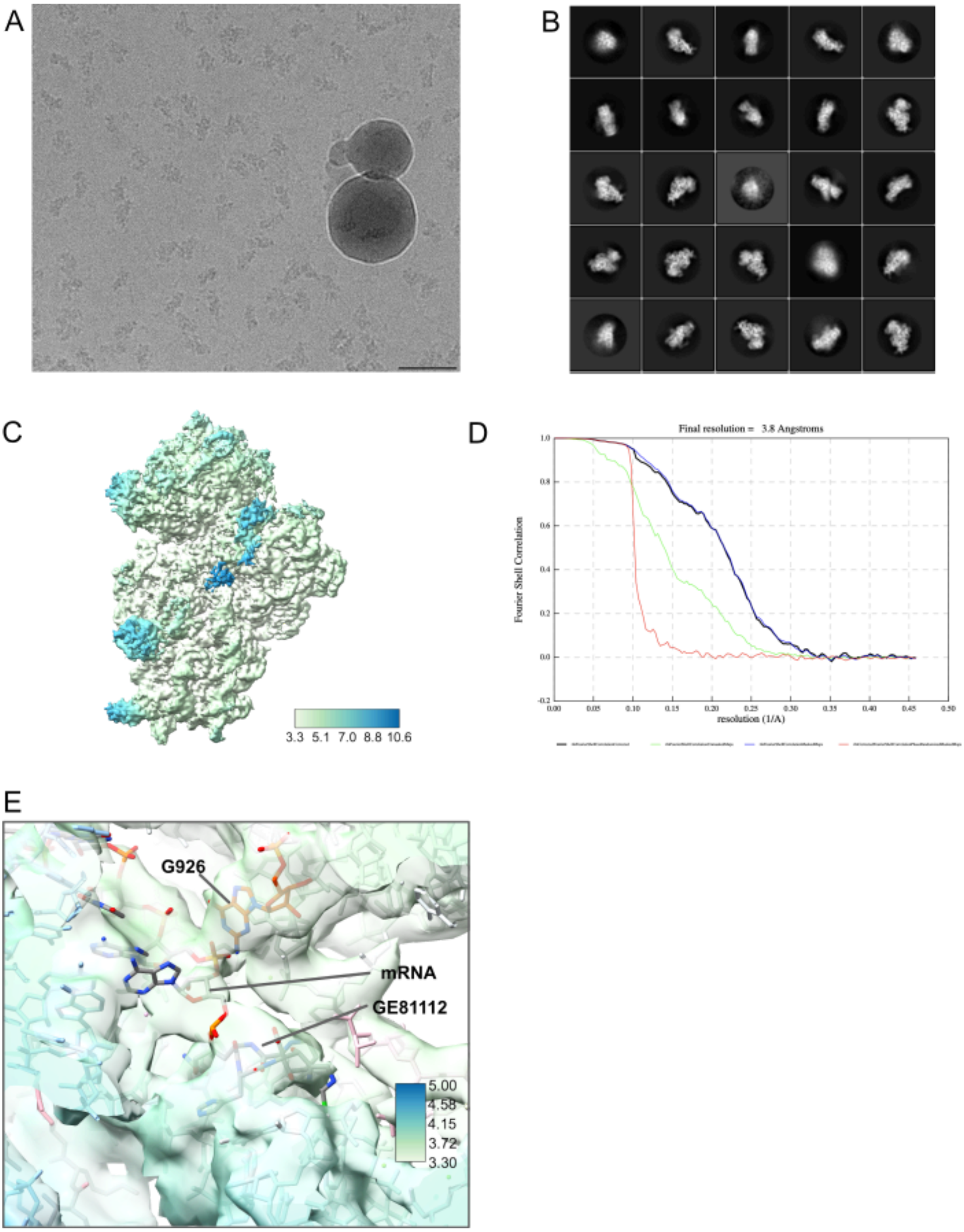


**Supplemental Figure S1: Cryo-EM Processing Results for Dataset 1.** (**A**) Representative cryoEM micrograph (**B**) 2D class averaging showing well-defined 30S ribosomal particles. (**C**) The final unsharpened cryoEM map (complex 1) is coloured according to local resolution (RELION). (**D**) The FSC curve corresponding to the cryoEM map seen in panel **C**. (**E**) The cryoEM map (unsharpened) surrounding the GE81112 binding site is coloured according to the local resolution estimate. GE81112, 16S rRNA residue G926 and the -1 mRNA residue are shown to highlight that in the lower resolution cryoEM map, the mRNA approaches GE81112. Note for the local resolution colouring, the max value was set to 5 Å; therefore, the dark blue represents 5 Å or greater.

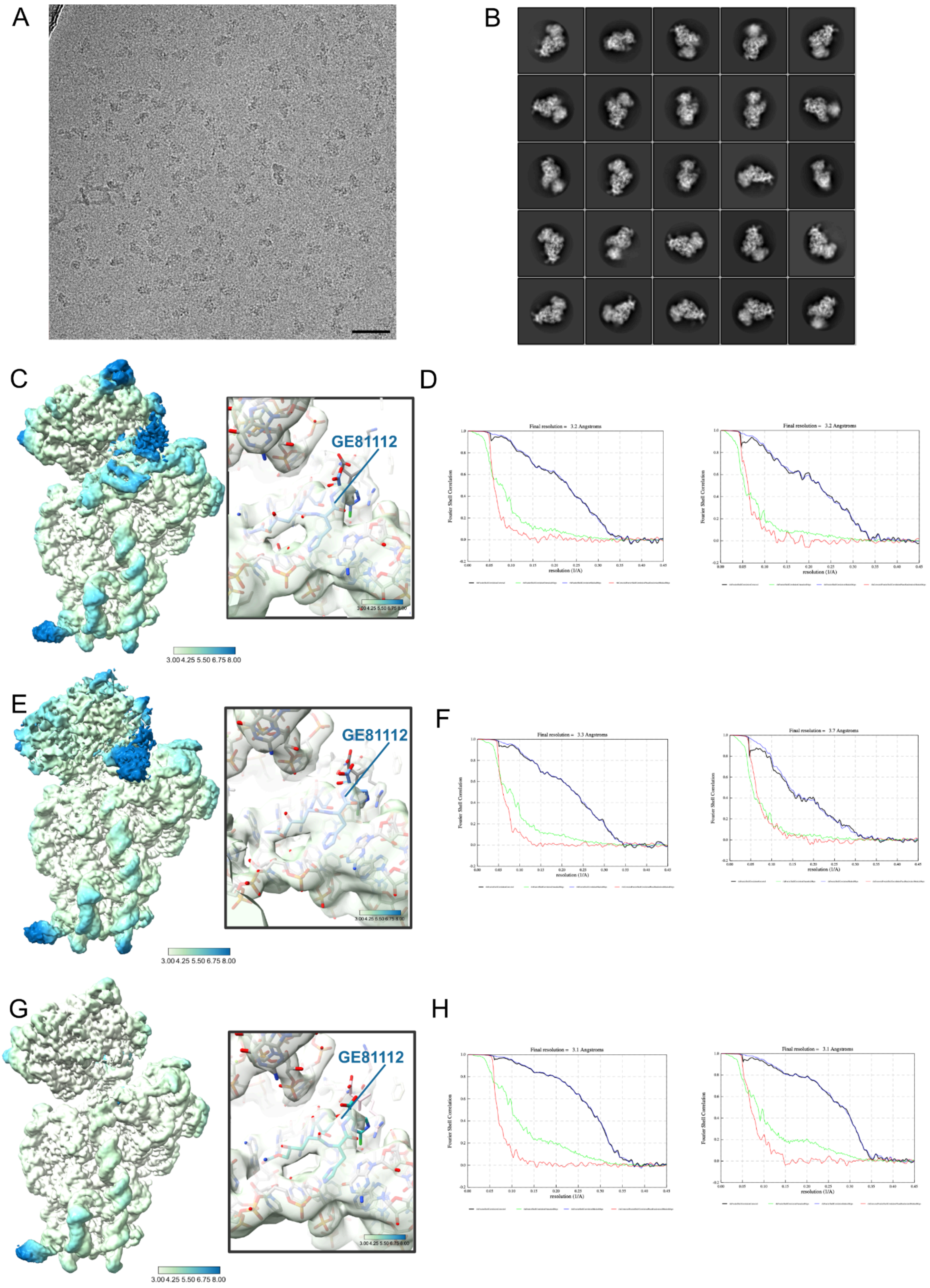


**Supplemental Figure S2: Cryo-EM Processing Results for Dataset 2.** (**A**) Representative cryoEM micrograph. The scalebar is 50 nm (**B**) Representative 2D classes. (**C**) Final multibody volumes from complex 2 colored by the local resolution (**D**) FSC curves for the multibody volumes in panel **C**; left, body; right head. (**E**) Final multibody volumes from complex 3 coloured by the local resolution (**F**) FSC curves for the multibody volumes in panel **C**; left, body; right head. (**G**) Final multibody volumes from complex 4 coloured by the local resolution (**H**) FSC curves for the multibody volumes in panel **C**; left, body; right head. The local resolution and fall-off in the FSC curves highlight the good quality of Complex 4 (**G** and **H**), while the low local resolution and bumpy FSC curve (**F**, **right**) indicates the body 2 map (30S head) of Complex 3 (**E** and **F**) is the poorest map from dataset 2. Note for the local resolution colouring, the max value was set to 8 Å; therefore, the dark blue represents 8 Å or greater.

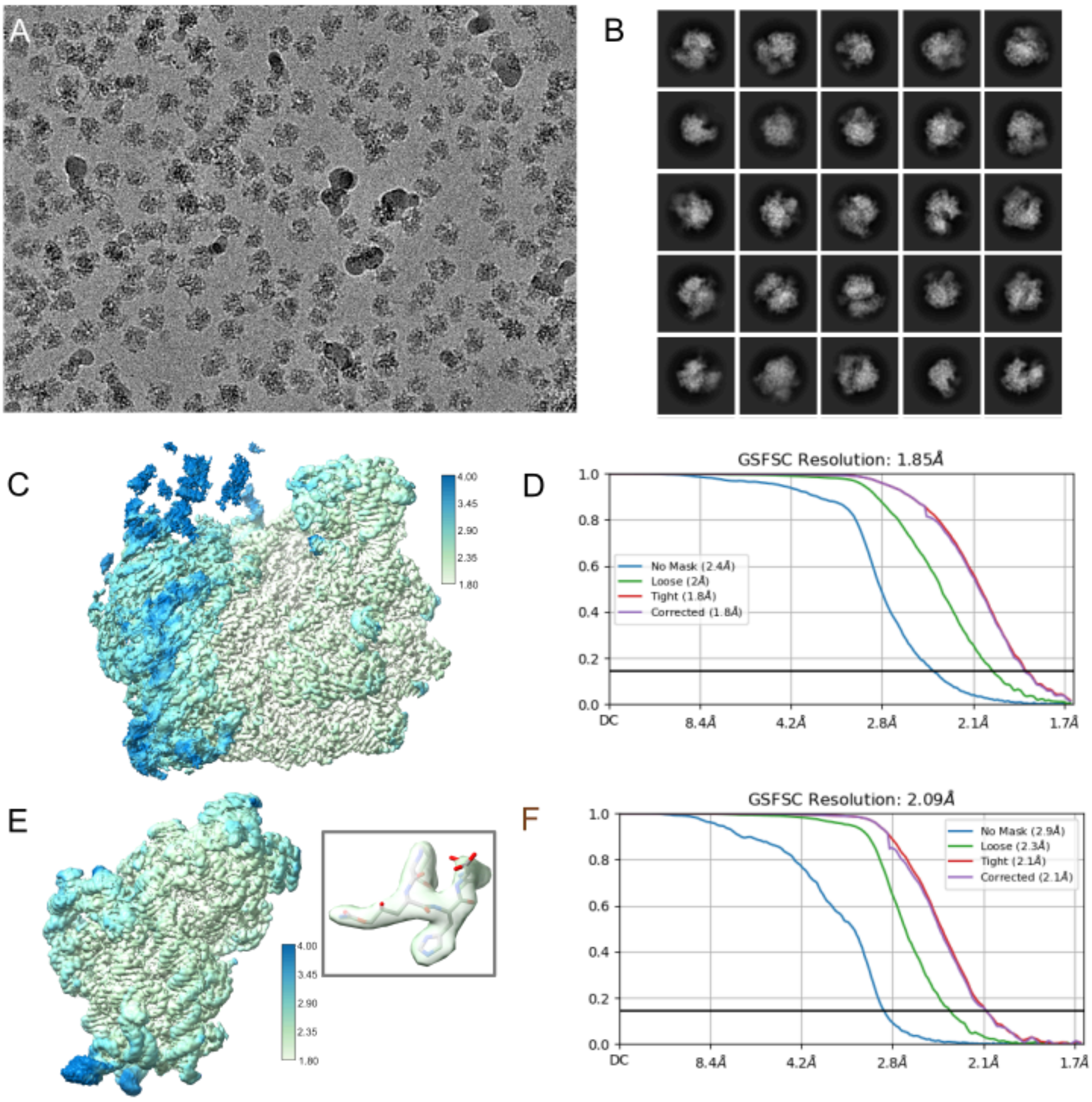


**Supplemental Figure S3: Cryo-EM Processing Results for Dataset 3.** (**A**) Representative cryoEM micrograph (**B**) 2D class averaging showing well-defined 70S ribosomal particles. (**C**) The unshapen cryoEM coloured according to local resolution (cryoSPARC) corresponds to a 70S particle refined under a mask for the 50S subunit. This volume was used in the subtraction job to yield projections of the 30S body that produced the cryoEM map seen in panel **E**. (**D**) The FSC curve corresponding to the cryoEM map seen in panel **C**. (**E**) The final cryoEM map (complex 5) corresponding to the 30S body region which harbours the GE81112 binding site (inset). The map is coloured according to the local resolution estimate (cryoSPARC) and masked to remove density for the 50S subunit that remained after subtraction. (**F**) The FSC curve corresponding to the cryoEM map seen in panel **E**. Note for the local resolution colouring, the max value was set to 4 Å; therefore, the dark blue represents 4 Å or greater.

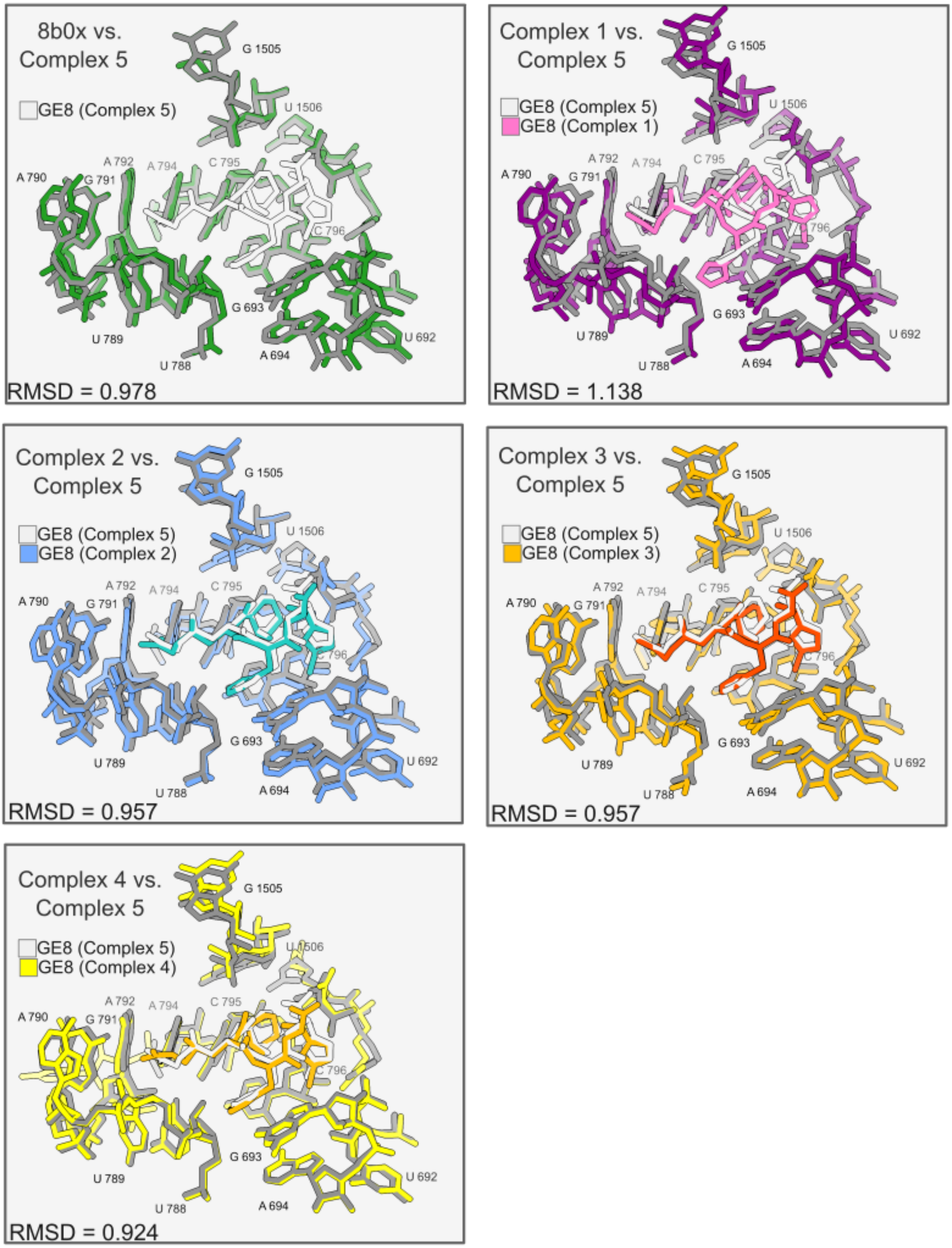
8b0x vs.
Complex 5
GE8 (Complex 5)
G 1505
U 1506
A 792
A 794
C 795
A 790
G 791
C 796
G 693
U 789
U 788
A 694
U 692
RMSD = 0.978
Complex 1 vs.
Complex 5
GE8 (Complex 5)
GE8 (Complex 1)
RMSD = 1.138
Complex 2 vs.
Complex 5
GE8 (Complex 5)
GE8 (Complex 2)
RMSD = 0.957
Complex 3 vs.
Complex 5
GE8 (Complex 5)
GE8 (Complex 3)
RMSD = 0.957
Complex 4 vs.
Complex 5
GE8 (Complex 5)
GE8 (Complex 4)
RMSD = 0.924

**Supplemental Figure S4: Comparison of GE81112 pocket in Complexes 1-5.** The models for the five complexes and a 70S ribosome bound by a P-tRNA (8b0x; (Fromm et al., 2023)) were aligned using residues within 15 Å of GE81112 and the root-mean-square deviation (RMSD) between two sets of atoms constituting the GE81112 binding pocket (16S rRNA: 692-694,788-796,1505,1506) was calculated with UCSF ChimeraX (Meng et al., 2023). Calculates values are: 0.978 (8b0x vs complex 5); 1.138 (complex 1 vs complex 5); 0.957(complex 2 vs complex 5); 0.957(complex 3 vs complex 5); 0.924(complex 4 vs complex 5).

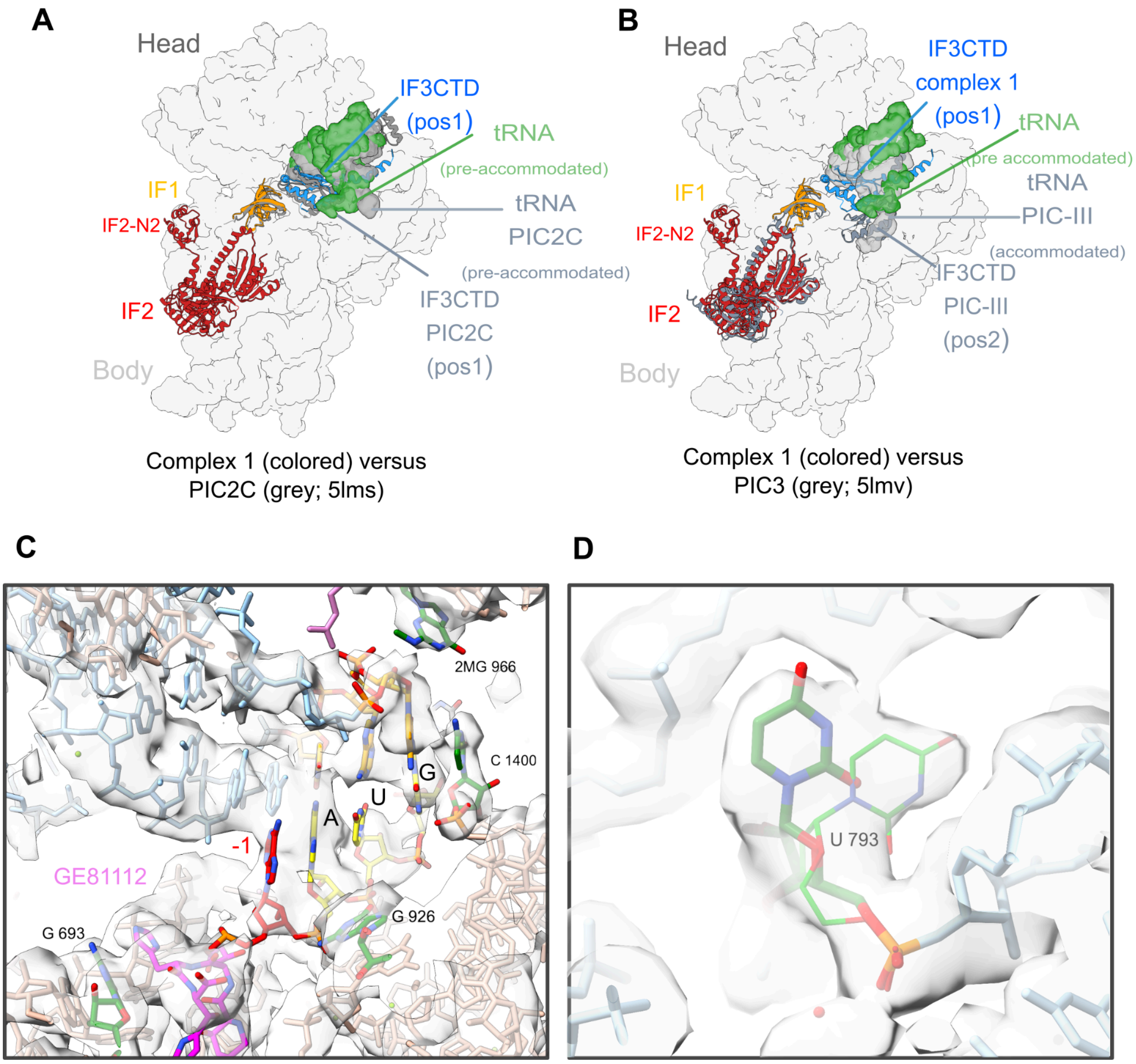


**Supplemental Figure S5: The initiation complex (Complex 1) are in a preIC state.** The position of fMet-tRNA, IF-1(orange), IF-2(crimson), and IF-3 in complex 1 are compared to that seen in the (**A**) PIC2C (5lms) and (**B**) PIC-III (5lmv) complexes of Hussain *et al.* (Hussain et al., 2016). The 30S body and head are shown as a flat surface and based on the complex 1 model. The alignment was made using 16S rRNA residues in the body domain. PIC2C and PIC-III show the IF3-CTD domain in 2 different positions, pos1 and pos2. Previously, we observed the GE81112 binding is associated with IF3-CTD being bound in pos1 (López-Alonso et al., 2017). As seen in panel **A,** the CTD of IF3 in complex I is positioned in pos1, and the fMet-tRNA is in a pre-accommodated position, indicating that GE81112 has trapped the complex in a preIC state. As seen in panel (**C**), an inspection of the codon-anticodon region of complex 1 is also indicative of the fMet-tRNA being in a pre-accommodated state, for example, the density for the +1 (U) basepair is weak, and the spacing between C1400 and G966 is too far for them to interact (5 Å) although C1400 and G966 are positioned to press against the last basepair of the codon-anticodon duplex. The backbone of the -1 residue shows some density, but the nucleobase is largely unaccounted for. Note we also observe density for the N2 domain of IF2 interacting near h16 (panels **A** and **B**), similar to that reported for the *Pseudomonas aeruginosa* initiation complex(Basu et al., 2022). (**D**) The high-resolution Complex 5 map (unsharpended) shows density indicative of an alternative confirmation of U793 (main conformation green thick sticks, alternative conformation thin green sticks). The maps of Complexes 1-4 show U793 in the main conformation. All chains other than the initiation factors have been rendered as a surface. In panels A and B, Complex 1 was used to visualise the IFs and fMet-tRNA, as in Complexes 2-4, we used MultiBody refinements and treated the body + IFs as a separate body from the head +tRNA/mRNA. As such, the models of the two bodies are out of alignment with each other after the refinement and density at the interface is problematic to interpret.

A

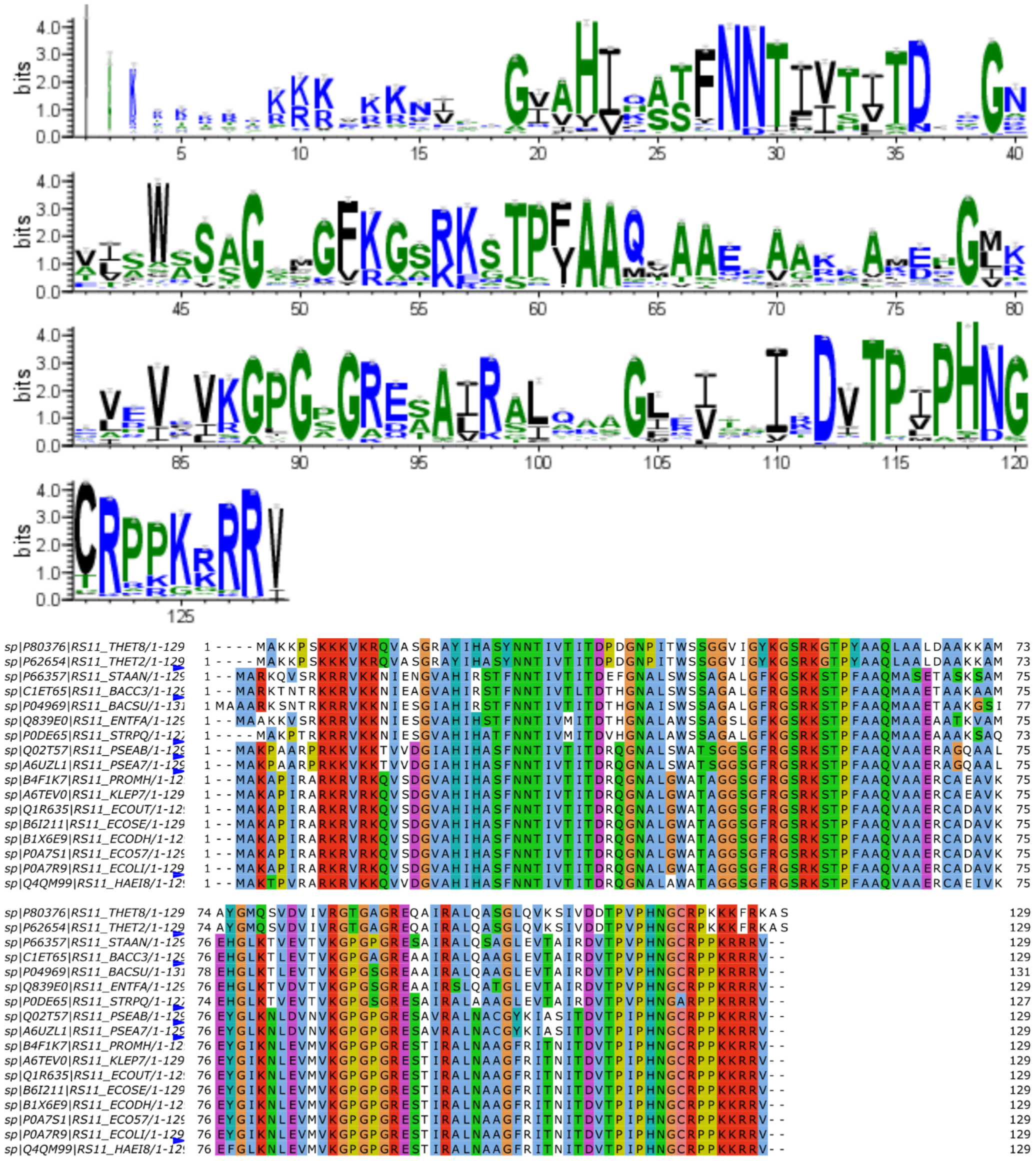

B

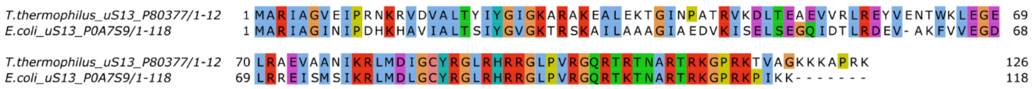

**Supplemental Figure S6: Conservation of r-proteins S11 and S13. (A)** The conservation of r-protein S11 is illustrated with WebLogo (top) with selected sequences aligned in below. (**B**) Clustal alignment of *T. thermophilus* (P80377) and *E. coli* (P0A7S9) r-protein S13.

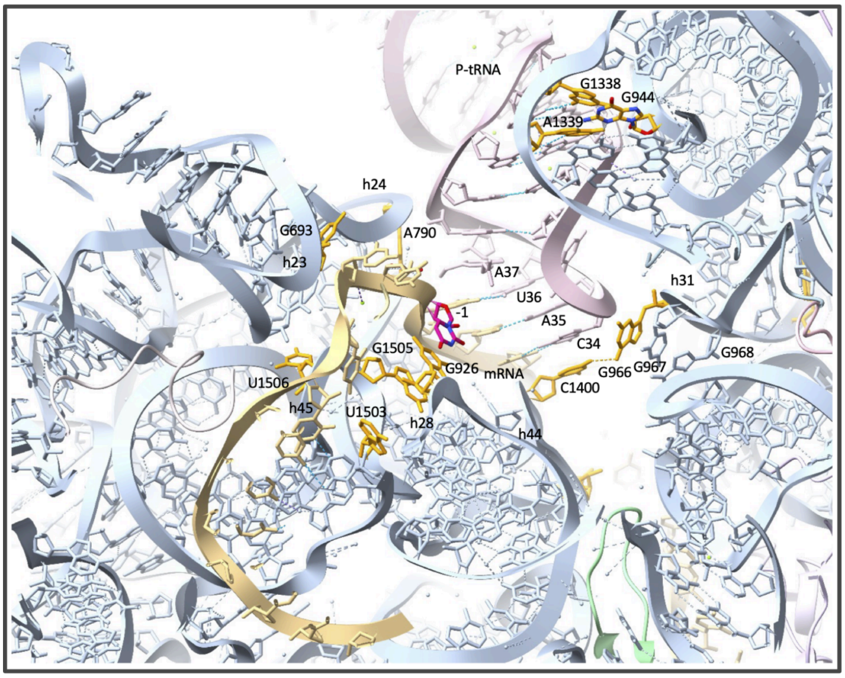
P-tRNA
G1338
G944
A1339
h24
G693
A790
h23
A37
U36
h31
-1
A35
C34
G1505
G968
G926
mRNA
G966
G967
U1506
C1400
h45
U1503
h28
h44

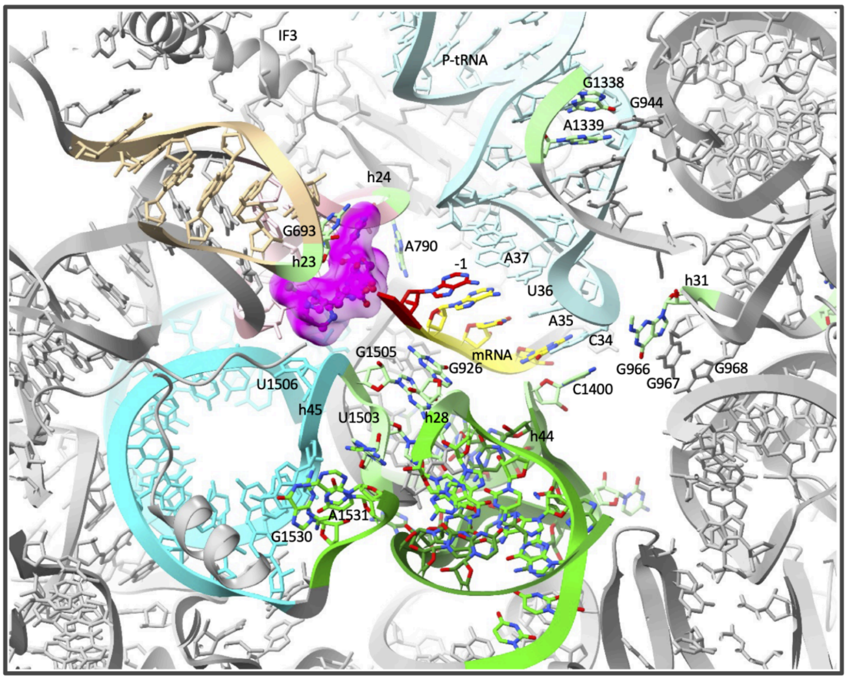
IF3
P-tRNA
G1338
G944
A1339
h24
G693
A790
h23
A37
-1
U36
h31
A35
C34
G1505
mRNA
G926
G966
G968
G967
U1506
C1400
h45
U1503
h28
h44
A1531
G1530

**Supplemental Figure S7: Key mRNA, tRNA and 30S interactions are altered or disrupted in the GE81112 pre-initiation complex.** GE81112 impedes the correct positioning of the mRNA (gold ribbon; top) in the E-site, hindering the formation of the interactions that characterise the full accommodation of the mRNA and tRNAs (depicted in top panel, 4v6f). Interestingly, these are formed between nucleotides located either at the tip/loop or in bulge positions of helices of the 30S body (i.e. G693,h23 tip (E-site mRNA and tRNA); 1503, 1505 and 1506, loop connecting h44 and h45 (E-site); 1400, h44 tip (P-sitetRNA); the universally conserved residues involved in the decoding and interacting with both mRNA and tRNA at the A-site: A1492 and A1493 in h44 and G530, in the "530 loop" of h18) and the head ((i.e. G926 in h28 (E-site, mRNA -1); C1054, in bulge position in h34 (mRNA and tRNA A-site, decoding) and 1534, which is part of the SD-aSD helix and interacts with 929 of h28; G1338 and A1339 in the loop that connects h29 to h42, (P-site tRNA); A790, tip h24, (P-site tRNA); G966, tip h31 (P-tRNA); G926, bulge position in h28, interacts with the mRNA P-site) (Mohan et al., 2014; Noller et al., 2017; Nishima et al., 2022; Hoffer et al.2020; Noller et al. 2022; Nguyen et al., 2023). Importantly, the ribosome does not interact with the mRNA at the P-site so that its positioning on this site is ensured through other interactions. In this respect, it could play a critical role the set of interactions between the ribosome and the mRNA at the E-site, represented by a strong polar hydrogen bond with the phosphate at the +1 mRNA position and a stacking interaction with the base of G926 with the mRNA in position -1 (Noller et al. 2022), depicted in red in the top panel. In the pre-initiation complex obtained in this study, in the presence of GE81112 (Complex 1, bottom panel), although G926 is involved in an interaction with a backbone phosphate of the mRNA, the base of the preceding mRNA nucleotide (depicted in red), does not perform a stacking interaction with G926 as it is instead base paired with A37 of the P-tRNA. Interestingly, in this complex, 3 additional bases of the mRNA are base paired with the tRNA, at position C34, A35 and U35. Interestingly, in this complex, 3 additional bases of the mRNA are base paired with the tRNA, at position C34, A35 and U35. In this complex, the residues described above, involved in the key interactions between mRNA, tRNA and the ribosome result in slightly different relative positioning, in concomitance with a different orientation of the head. In this respect, the key interaction between G966 and C1400 that indeed stabilises the closed conformation of the head of the ribosome, is not performed as the two residues result at a distance of more than 6A.
